# Coupling effect of morphology and mechanical properties contributes to the tribological behaviors of snake scales

**DOI:** 10.1101/148734

**Authors:** Long Zheng, Yinghui Zhong, Yihang Gao, Jiayi Li, Zhihui Zhang, Zhenning Liu, Luquan Ren

## Abstract

It is known that the tribological behaviors of snake skins are contributed by the synergistic action of multiple factors, such as surface morphology and mechanical properties, which has inspired fabrication of scale-like surface textures in recent years. However, the coupling effect and mechanism remain to be elucidated. In this work, the morphology and mechanical properties of the scales from different body sections (leading body half, middle trunk and trailing body half) and positions (dorsal, lateral and ventral) of *Boa constrictor* and *Eryx tataricus* have been characterized and compared to investigate the corresponding effects on the tribological behaviors and to probe the possible coupling mechanism. The morphological characterizations of scanning electron microscopy and atomic force microscopy have revealed significant differences between the two species with the roughness of scales from *Boa constrictor* being larger in general. The mechanical properties measured by nanoindentation have corroboratively demonstrated substantial differences in terms of elastic modulus and hardness. Meanwhile, tribological characterizations of scales in different body positions from the two species also exhibit evident anisotropy. Interestingly, the ventral scales manifest higher friction coefficients but lower surface roughness, together with relatively larger elastic modulus and hardness. A “double-crossed” hypothesis has been proposed to explain the observed coupling effect of the morphology and mechanical properties on friction, which may afford valuable insights for the design of materials with desirable tribological performance.

## 1. Introduction

Evolution and selection in nature has yielded various species with optimized biological features, which have inspired bionic engineering over recent decades (Ball, 1999; Comanns et al., 2016; Huang et al., 2012; Meyers et al., 2008; Wong et al., 2011). Snakes are one of nature’s most amazing well-adapted reptilians without extremities, which have evolved over 150 million years and inhabited every land on earth except the coldest regions (Evans, 2003; Klein and Gorb, 2014). One interesting trait of snakes is the sliding locomotion, which renders the ventral scales at the belly in direct and continuous contact with the surroundings. Therefore, the snake scales have developed a range of tribological characteristics fitting to respective habitat conditions, which could afford valuable insights for friction and wear research (Abdel-Aal et al., 2012).

Friction remains a big concern in engineering and recently, more investigations have been made on the structures and materials of snake scales to achieve better understanding of the frictional behaviors (Abdel-Aal et al., 2012; Arzt et al., 2003; Benz et al., 2012; Berthé et al., 2009; Hazel et al., 1999; Klein et al., 2010; Klein and Gorb, 2012; Marvi and Hu, 2012; Rechenberg, 2003; Rocha-Barbosa and Moraes e Silva, 2009). In these reported cases, snake scales have manifested various desirable performance, such as adhesion reduction (Arzt et al., 2003), wear resistance (Rocha-Barbosa and Moraes e Silva, 2009) and frictional anisotropy (Benz et al., 2012). In particular, it has been suggested that distinct skin microstructures account for different frictional properties in discrete body positions (Berthé et al., 2009). It has also been shown that the functional microstructure on scales is an adaptation of snake species to their preferential habitats (Rocha-Barbosa and Moraes e Silva, 2009). Moreover, mechanical analysis of snake epidermis has revealed a gradient of hardness and elastic modulus, which corresponds to the respective inhabiting conditions, indicating that such a gradient is a potential adaptation to locomotion and wear reduction on substrates (Klein et al., 2010; Klein and Gorb, 2012).

The insights gained from the investigations of snake epidermis have inspired various design mimicking natural systems (Abdel-Aal and El Mansori, 2011; Abdel-Aal and El Mansori, 2013; Baum et al., 2014a; Baum et al., 2014b; Cuervo et al., 2016; Greiner and Schäfer, 2015; Mühlberger et al., 2015). For instance, it has been suggested that a surface texture resembling the scale microstructures of the *Python regius* is able to benefit the lubrication of cylinder liners (Abdel-Aal and El Mansori, 2011). The frictional examination on polymer surface with snake-inspired microstructures has revealed that the special ventral surface ornamentation of snakes cannot only lower friction coefficient, but also generate anisotropic friction, which might be another adaptation to sliding locomotion (Baum et al., 2014a; Baum et al., 2014b).

These works have also indicated that the frictional behaviors of snake skins are contributed by the synergistic actions or coupling effects of multiple factors rather than determined by a single factor (Ren, 2009; Zheng et al., 2016). Yet, the coordinated mechanism of multiple factors remains to be elucidated in order to implement bio-mimicking tribosystems with desirable friction performance.

The objective of the present study is to investigate the effects of morphology and mechanical properties on the frictional behaviors of snake epidermis and to probe the coupling effect between these two factors as a result of possible adaptation. The following three key questions are addressed in this work: (1) What are the major morphological differences between the two surveyed snake species (*Eryx tataricus* and *Boa constrictor*) from different inhabiting conditions, especially with regards to distinct body positions? (2) How do the mechanical properties of scales vary between these two species? (3) How does the coupling of surface roughness and mechanical properties take effect on the frictional behaviors? The shed snake epidermis from *Eryx tataricus* (Squamata) and *Boa constrictor* (Squamata) from different preferential habitats and thus moving on different substrates has been compared to address these questions.

## 2. Materials and methods

### 2.1 Animals

Two snake species, *Boa constrictor* and *Eryx tataricus*, from distinct preferential habitats were chosen for the study.

*Boa constrictor* (Linnaeus, 1758), also called red-tailed boa, is a species of large, heavy-bodied snake in the *Boidae* family. *Boa constrictor* lives in a broad range of environments, from tropical rainforests to arid semidesert country (Stidworthy, 1974). However, it is more often found in rainforests due to the preferred humidity and temperature as well as plenty of potential preys. As semi-arboreal snakes, young *Boa constrictors* may climb onto trees and shrubs to fodder. However, they become mostly terrestrial as they grow older and heavier (Mehrtens, 1987).

*Eryx tataricus*, commonly found from the east Caspian Sea to China west and Mongolia, is a non-venomous snake species in the *Boidae* family, which was first described by Lichtenstein in 1823 (Wallach et al., 2014). It normally burrows under sands or rocks via the locomotion of lateral undulation at a relative slow speed. The inhabiting conditions of *Eryx tataricus* are arid and warm.

The animals were reared in 80 cm × 60 cm × 60 cm (length × width × height) wood cabinets under controlled temperature and humidity at Jilin University, China. The bottom of the cabinet for *Eryx tataricus* was covered by a sand layer of roughly 10 cm, whereas no sand was added to the other cabinet for *Boa constrictor*. Two adult mice were fed twice a week. All samples examined in this study were shed skins derived from a 5 year-old male *Eryx tataricus* (body mass of 0.25 kg and length of 115 cm, Fig. 1, top) and a 3 year-old male *Boa constrictor* (body mass of 1.0 kg and length of 193 cm, Fig. 1, bottom).

**Figure 1.**
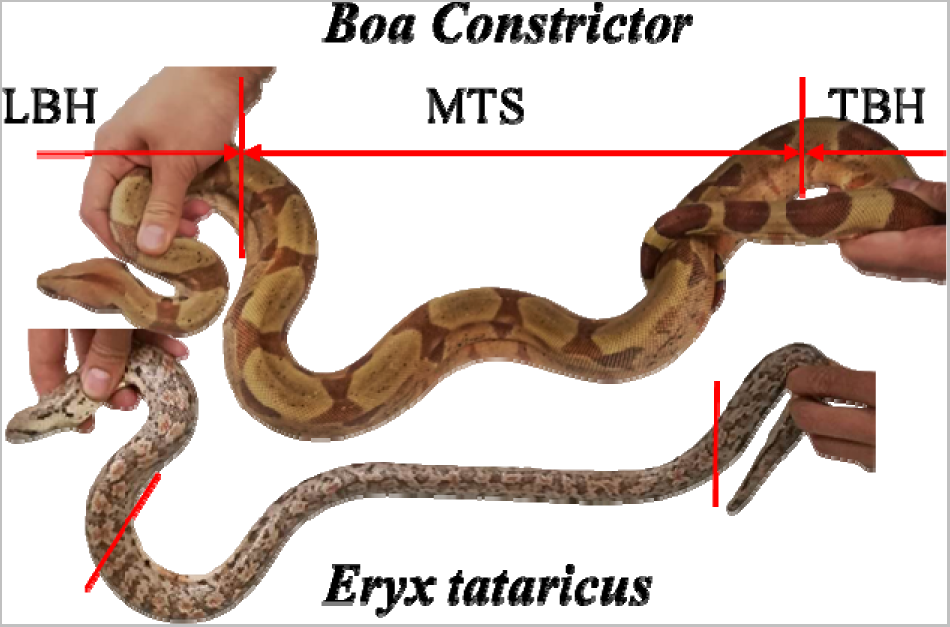
The photographs of *Boa Constrictor* and *Eryx tataricus*. LBH: Leading Body Half; MTS: Middle Trunk Section; TBH: Trailing body Half.

To characterize the morphological features of the shed epidermis from different body positions (dorsal, lateral and ventral), the snakes are divided into three major sections (Fig. 1) as previously described (Hisham, 2013). The first one is denoted as Leading Body Half (LBH) near the head. The stockiest portion of the body is regarded as Middle Trunk Section (MTS), which follows LBH as the second section. The remaining portion is named as Trailing body Half (TBH), which accounts for approximately the same length ratio as LBH but near the tail.

### 2.2 Sample collection and treatments

The epidermis of adult individuals was collected directly after shedding from the cabinets and washed repeatedly to remove excreta, followed by soaking in distilled water for four hours at room temperature to unfold the wrinkles. Subsequently the shed epidermis was wiped with paper towels. Then the shed skins were dried by a hair dryer and wrapped in baking paper, which were then stored in sealed plastic bags at room temperature until use.

### 2.3 Scanning electron microscopy (SEM)

The samples of dorsal, lateral and ventral scales from three sections of the two species were cut into 5 mm ×5 mm pieces and mounted on microscope slides with cyanoacrylate adhesive (Ruichang Gold 3-Second Ltd. Co., China). The mounted samples were sputter coated with gold-palladium of 10 nm in thickness and then observed by a scanning electron microscope (Zeiss EVO 18, England, Cambridge) at an accelerating voltage of 20 kV.

### 2.4 Atomic force microscopy (AFM)

The surface profiles of the samples were obtained by atomic force microscopy (AFM) (Bruker Dimension ICON, America) under the ScanAsyst-air contact mode. Scans were carried out at room temperature with a scan rate of 1 Hz and a resolution of 256 × 256 pixels using a ScanAsyst-air cantilever. The width, depth and height between two micro-units were estimated from AFM images processed by NanoScope analysis (Bruker GmbH, Germany version 1.40). The average roughness of samples were calculated with the same software.

### 2.5 Nanoindentation

The nanoindentation measurements were performed on a Nanoindenter (Agilent, G200) equipped with a Berkovich tip at a drift rate of 0.1 nm·s^-1^. All measurements were conducted with Poisson ratio of 0.3 and peak hold time of 50 s at room temperature. Herein, two different modes were used. (1) Under the basic mode, the elastic moduli and hardnesses of the samples can be calculated from the force and displacement curves. The elastic moduli and hardnesses of scales (5×5 mm) were examined by following the line of circles from the anterior edge to the posterior edge with a spacing of 300 μm, as depicted in Fig. 2A. At least 5 independent indentations were carried out at each distance denoted from the front with a spacing of 50 μm (Fig. 2A). (2) The CSM mode was applied on the ventral scales of *Boa constrictor* with a penetration depth up to 12000 nm (Fig. 2B) to acquire the profiles of elastic modulus and hardness with increasing penetration depth.

**Figure 2.**
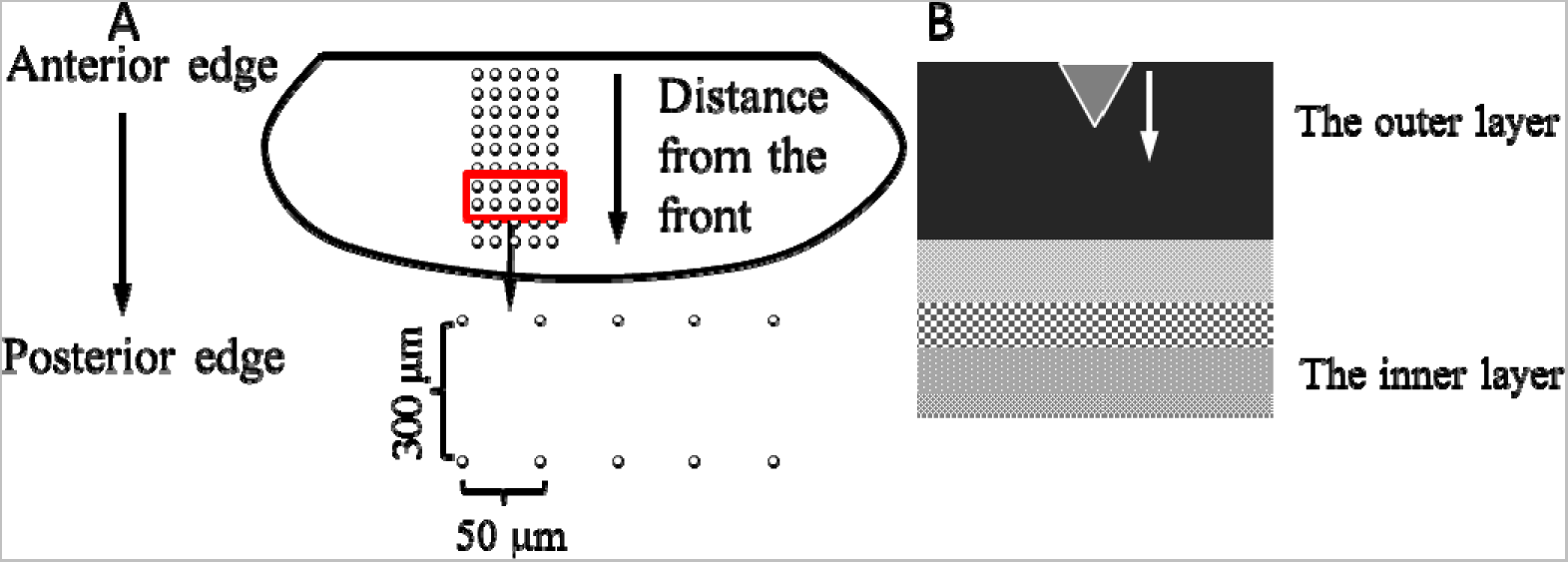
Schematic illustration of nanoindentation measurements. (A) The indented points on a single scale with circles representing test points. (B) The illustration of a probe penetrating the scale vertically to measure elastic modulus and hardness with increasing penetration depth. The inverted triangle represents the probe.

### 2.6 Tribological measurements

Dynamic tribological measurements were conducted on a nanotribometer (NTR^2^, Anton Paar Company, Switzerland) under the linear reciprocating mode with a maximum linear speed of 63 μm·s^-1^ and a stroke of 1 mm. The applied normal force was 0.6 mN. The samples were tested against a smooth, spherical silicon carbide probe (diameter≈2 mm) under a dry-sliding contact condition at room temperature (RH≈40%). All samples were mounted on microscope slides using cyanoacrylate adhesive. The samples were tested in the forward, backward and transverse directions. For each kind of scale, six parallel measurements were carried out on triplicate scales with duplicated measurements on each. The friction coefficients were recorded by the software (version 6.2.10) and statistically analyzed with Origin (version 8.5) and SPSS (version 22.0) (Students’ t test).

## 3. Results

### 3.1 Morphology on scale surface

Previous work has demonstrated that scale microstructures play important roles in the tribological properties of snake skins (Abdel-Aal et al., 2010; Abdel-Aal, 2016; Klein and Gorb, 2016). Hence, we first set out to examine the microstructures of shed scales from *Boa constrictor* and *Eryx tataricus* by scanning electron microscopy (SEM) (Fig. 3). Generally speaking, the microstructures on lateral scales from middle-trunk section (MTS) of *Boa constrictor* (Fig. 3B) and *Eryx tataricus* (Fig. 3E) exhibit similar patterns of roughly longitudinal-aligned flakes with overlapping-caused ridges, whereas the dorsal (Fig. 3A and 3D) and ventral (Fig. 3C and 3F) scales reveal different micro-morphologies for two species. The surface patterns on the ventral scale of *Boa constrictor* (Fig. 3C) and the dorsal scale of *Eryx tataricus* (Fig. 3D) resemble the common feature observed for the lateral scales. In contrast, well-aligned ridges paralleled to the longitudinal body axis are found on the dorsal scales from the MTS of *Boa constrictor* (Fig. 3A). Moreover, the SEM image of the ventral scales from the MTS of *Eryx tataricus* display a relatively smooth morphology with no practically discernible micro-ornaments at the same magnification (Fig. 3F), which is in sharp contrast to those of the dorsal and lateral scales. Random scratches and wear patterns have been frequently found on these ventral scales (Fig. 3H). It is noteworthy that the front edges of the scales of *Eryx tataricus* are mounted by tiny wrinkles (insets in Fig. 3D, 3E and 3F), which differ from the major portions of scales (Fig. 3D, 3E and 3F) and look like aeolian sand ripples (Fig. 3I). The transition from ripple-like wrinkles to smooth surface is also verified, as shown in Fig. 3G.

**Figure 3.**
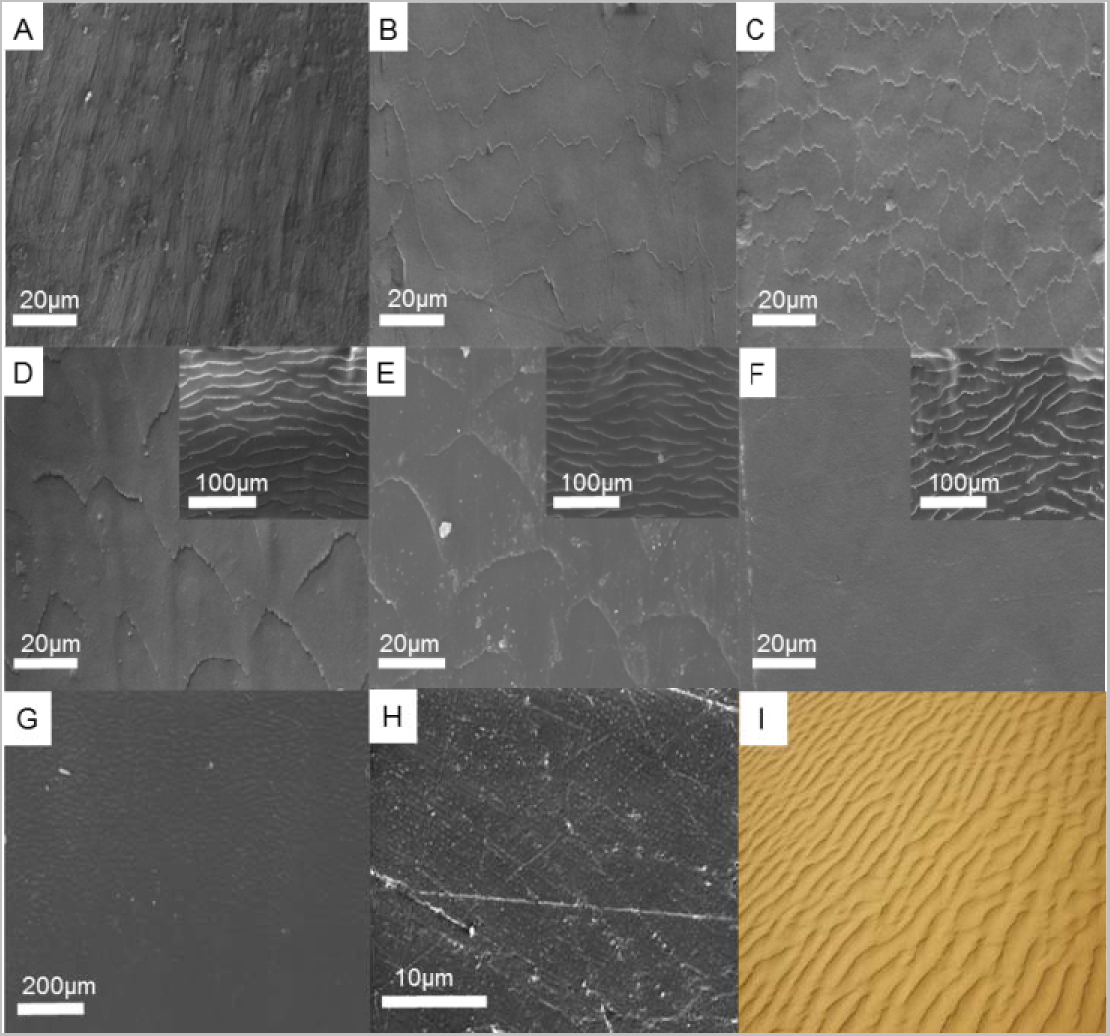
SEM images of shed scales (A-H) from two species and a photograph of aeolian sand ripple (I). (A), (B) and (C) show dorsal, lateral and ventral scales from the MTS of *Boa constrictor*. (D), (E) and (F) exhibit dorsal, lateral and ventral scales from the MTS of *Eryx tataricus*. The insets in (D), (E) and (F) are the microstructures on the front edges of respective scales of *Eryx tataricus*. (G) is an image of the transition from the front edge to the middle portion of the ventral scale of *Eryx tataricus*. Anterior is oriented at the top in image (A-G). (H) shows scratches and wear patterens on the vertral scale surface of *Eryx tataricus*. (I) is a photograph of aeolian sand ripple as a comparison (a courtesy from Sina website).

Surface roughness is one key parameter for morphological characterization and plays an important role in surface friction (Mühlberger et al., 2015; Voigt et al., 2012). Hence, the average surface roughness (Ra) of the scales from LBH, MTS and TBH of *Boa constrictor* and *Eryx tataricus* was measured by AFM (Fig. 4). Overall, the ventral scales of the two species demonstrate obviously lower Ra than the other body positions, albeit with microstructures on the scale surface. With regard to different body sections of the two species, the Ra of scales from LBH, MTS and TBH exhibit a consistent trend of dorsal > lateral > ventral with one exception at the LBH of *Eryx tataricus*, where the dorsal and lateral scales display comparable roughness. It is noteworthy that more significant difference of Ra has been observed for scales from *Eryx tataricus* in terms of either body positions or body sections. Moreover, the comparison between *Boa constrictor* and *Eryx tataricus* reveals a most prominent difference for dorsal scales at TBH.

**Figure 4.**
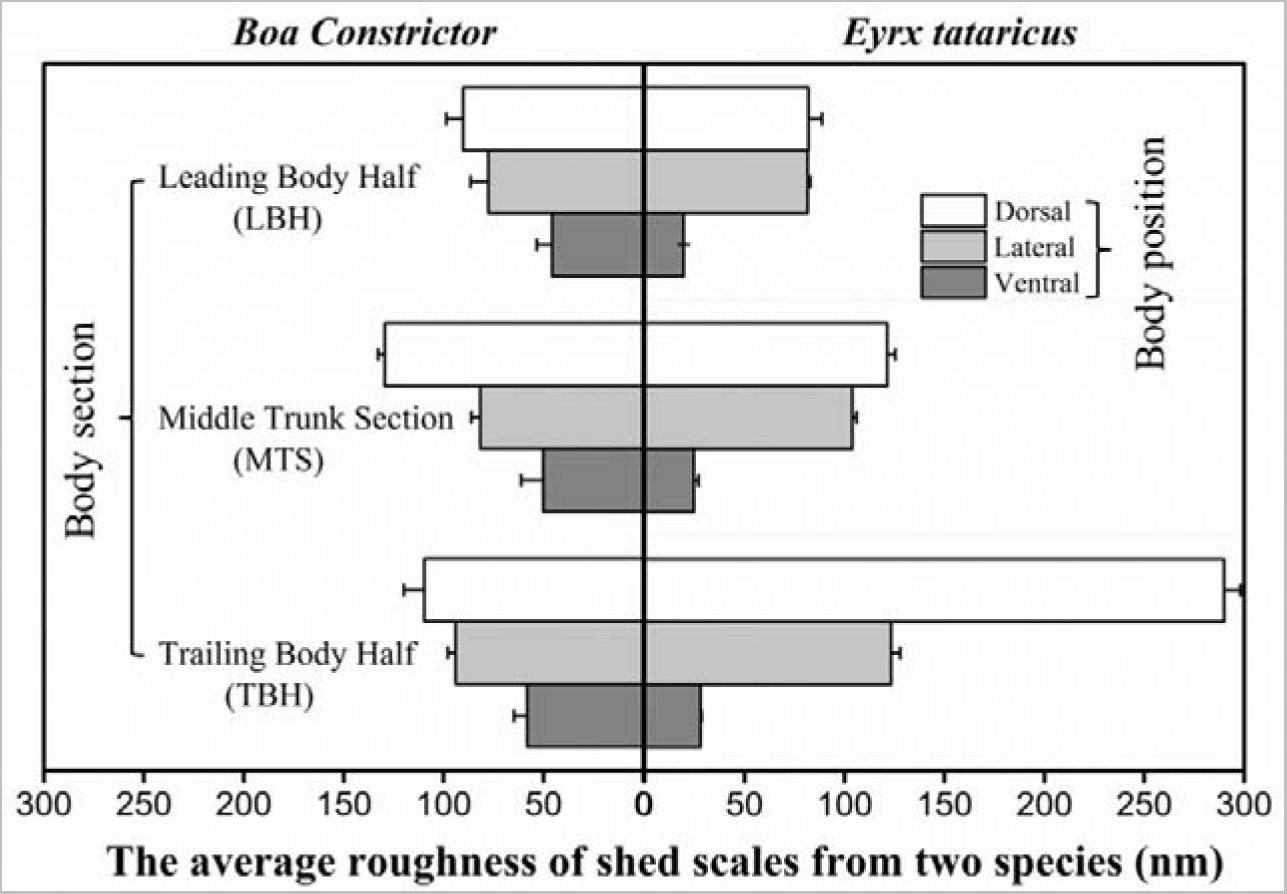
The average roughness (Ra) of scales in different body positions and sections of *Boa constrictor* and *Eryx tataricus*. The error bars denote standard deviations.

In order to gain a better understanding of the above trends, the parameters of unit microstructures, including the width, depth and height of micro-units, were extracted from AFM profiling and compared (Fig. 5 and Table 1). It is found that the widths, depths and heights of lateral scale units in different body sections of *Boa constrictor* are significantly larger than those of respective ventral scale units. In contrast, the widths and depths of dorsal scale units in different body sections of *Eryx tataricus* are close to those of respective lateral scale units, with one exception that the depth of dorsal scale units at TBH is more than three times of the depth of lateral scale units. In addition, the heights of dorsal scale units in different body sections of *Eryx tataricus* are also larger than those of lateral scale units. In particular, for TBH the height of dorsal scale units is approximately 6.5 times of the height of lateral scale unit. These parameters (Table 1) have corroborated the measured roughness (Fig. 4) in following aspects: first, for *Boa constrictor*, lateral scales are rougher than ventral scales regardless of body sections; second, for *Eryx tataricus*, the roughness of dorsal scales and lateral scales are comparable at LBH and MTS; third, the dorsal scales at TBH of *Eryx tataricus* show more dramatic micro-patterning and thus higher roughness. It should be noted that it is difficult to extract unit parameters for the dorsal scales of *Boa constrictor* and the ventral scales of *Eryx tataricus*.

**Figure 5.**
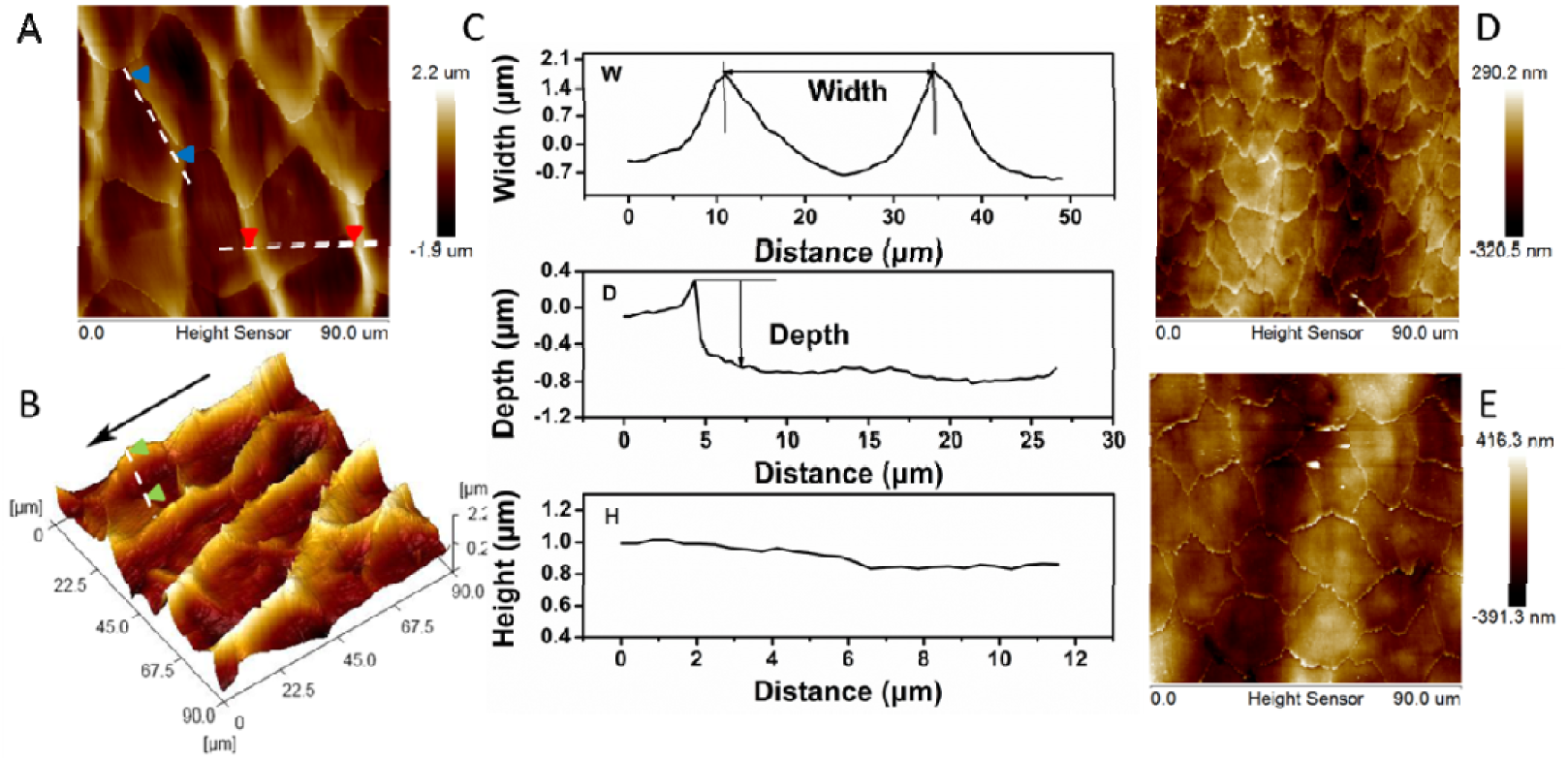
Representative AFM images (A, B, D and E) and parameter extractions from AFM profiling (C). (A) is an AFM image of microstructures on the dorsal scale of TBH from *Eryx tataricus* with indicated width profiling (between two red arrowheads) and height profiling (between two blue arrowheads). (B) is the 3D image of (A) with indicated depth profiling (between two green arrowheads). The black arrow on the side points to the posterior in (B). (C) shows the width (top), depth (middle) and height (bottom) profiles along the corresponding lines in (A) and (B). The width, depth and height extracted from (C) are 23.761 m, 0.989 m and 0.916 m, respectively. (D) and (E) are AFM images of microstructures on the ventral and lateral scales of MTS from *Boa constrictor*. W: the width between the two units of microstructures. D: the height difference between the ridge and the valley. H: the height measured along the ridge.

**Table 1.** Parameters for unit microstructures on scales from *Boa constrictor* and Eryx tataricus

| Scales | Width ($\mu\text{m}$) | Depth ($\mu\text{m}$) | Height ($\mu\text{m}$) |
| --- | --- | --- | --- |
| ***Boa constrictor*** |  |  |  |
| Ventral (LBH) | 13.362±1.647 | 0.128±0.010 | 0.036±0.008 |
| Ventral (MTS) | 13.221±0.640 | 0.139±0.011 | 0.060±0.009 |
| Ventral (TBH) | 18.557±0.786 | 0.138±0.015 | 0.042±0.005 |
| Lateral (LBH) | 20.727±0.967 | 0.173±0.015 | 0.073±0.012 |
| Lateral (MTS) | 20.509±0.858 | 0.258±0.017 | 0.082±0.010 |
| Lateral (TBH) | 25.457±1.097 | 0.203±0.012 | 0.055±0.011 |
| ***Eryx tataricus*** |  |  |  |
| Dorsal (LBH) | 25.536±1.850 | 0.321±0.017 | 0.144±0.011 |
| Dorsal (MTS) | 27.751±1.018 | 0.494±0.049 | 0.134±0.015 |
| Dorsal (TBH) | 23.229±1.067 | 0.992±0.114 | 1.036±0.125 |
| Lateral (LBH) | 28.610±1.945 | 0.344±0.037 | 0.104±0.016 |
| Lateral (MTS) | 28.022±0.961 | 0.451±0.036 | 0.127±0.014 |
| Lateral (TBH) | 26.908±1.124 | 0.281±0.055 | 0.159±0.017 |

### 3.2 Mechanical properties of scales

It has been reported that snake skins are equipped with anti-wear functions owing to their mechanical properties (Klein et al., 2010). To this end, nanoindentation was utilized to characterize the mechanical properties (*i.e.* elastic modulus and hardness) of scales in different body positions and sections from *Eryx tataricus* and *Boa constrictor* (Fig. 6-8). Fig. 6A exhibits elastic moduli and hardnesses of a single scale with different penetration depths in the vertical direction. Consistent with previous findings (Klein et al., 2010), the measured elastic moduli and hardnesses become relatively stable beyond the penetration depth of 200 nm. Then the elastic modulus displays a slight rise as the penetration increases, whereas the hardness slowly declines. Such a depth-correlated gradient has also been reported by other groups (Klein and Gorb, 2012). It is noted that the elastic modulus and hardness are almost constant at depth from 200 to 2000 nm (inset in Fig. 6A). Therefore, the elastic moduli and hardnesses at the depth of 1000 nm were measured from anterior to posterior on a single scale, as depicted in Fig. 2, to examine the horizontal changes of mechanical properties on scale surfaces. Fig. 6B and 6C show the elastic moduli and hardnesses of ventral scales from different body sections of *Eryx tataricus*, respectively. Interestingly, both elastic modulus and hardness are low near the front edge and gradually elevate to a steady level, indicating a horizontal gradient of mechanical properties also exists on the snake scale and the snake body as a whole can be regarded as an integration of alternating hard-soft materials.

**Figure 6.**
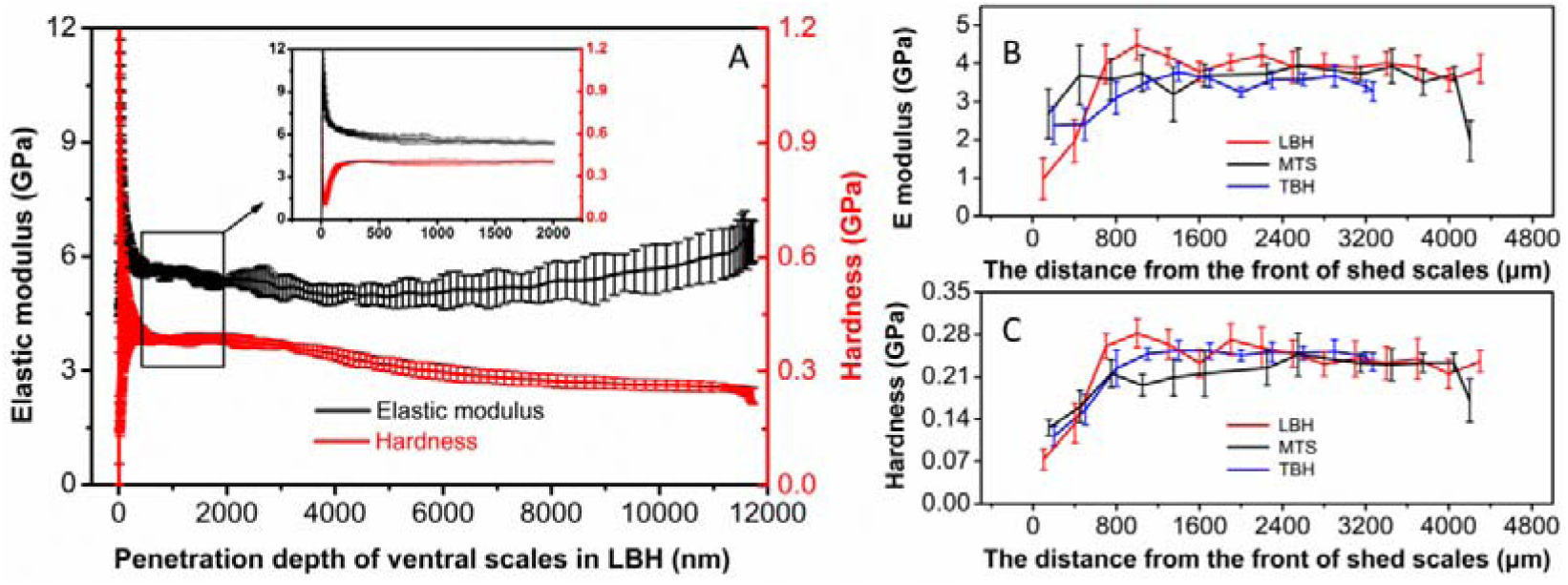
Elastic moduli and hardnesses of ventral scales in different sections of *Boa constrictor* and *Eryx tataricus*. (A) exhibits elastic moduli (black) and hardnesses (red) of ventral scales from *Boa constrictor* at different penetration depths. The inset shows elastic moduli and hardnesses of ventral scales at the depth of 200-2000 nm. (B) and (C) show elastic moduli and hardnesses from anterior to posterior on ventral scales in different sections of *Eryx tataricus*, respectively. The error bars denote standard deviations.

In order to compare the mechanical properties of scales across the whole body for the two species, nanoindentation (at penetration depth of 1000 nm) was carried out at the middle of scales from different body positions and sections, where the elastic modulus and hardness maintained a relatively stable level. Overall, the hardnesses of the two species are comparable, with those of *Boa constrictor* being a slightly higher (lines in Fig. 7). Yet, the scales of *Boa constrictor* exhibit generally larger elastic moduli than the counterpart of *Eryx tataricus* (histograms in Fig. 7). A common order of ventral > dorsal > lateral is also found for the elastic moduli and hardnesses of scales in different body sections of the two snakes, except that the hardness of dorsal scales at TBH of *Boa constrictor* is a bit smaller than that of respective lateral scales.

**Figure 7.**
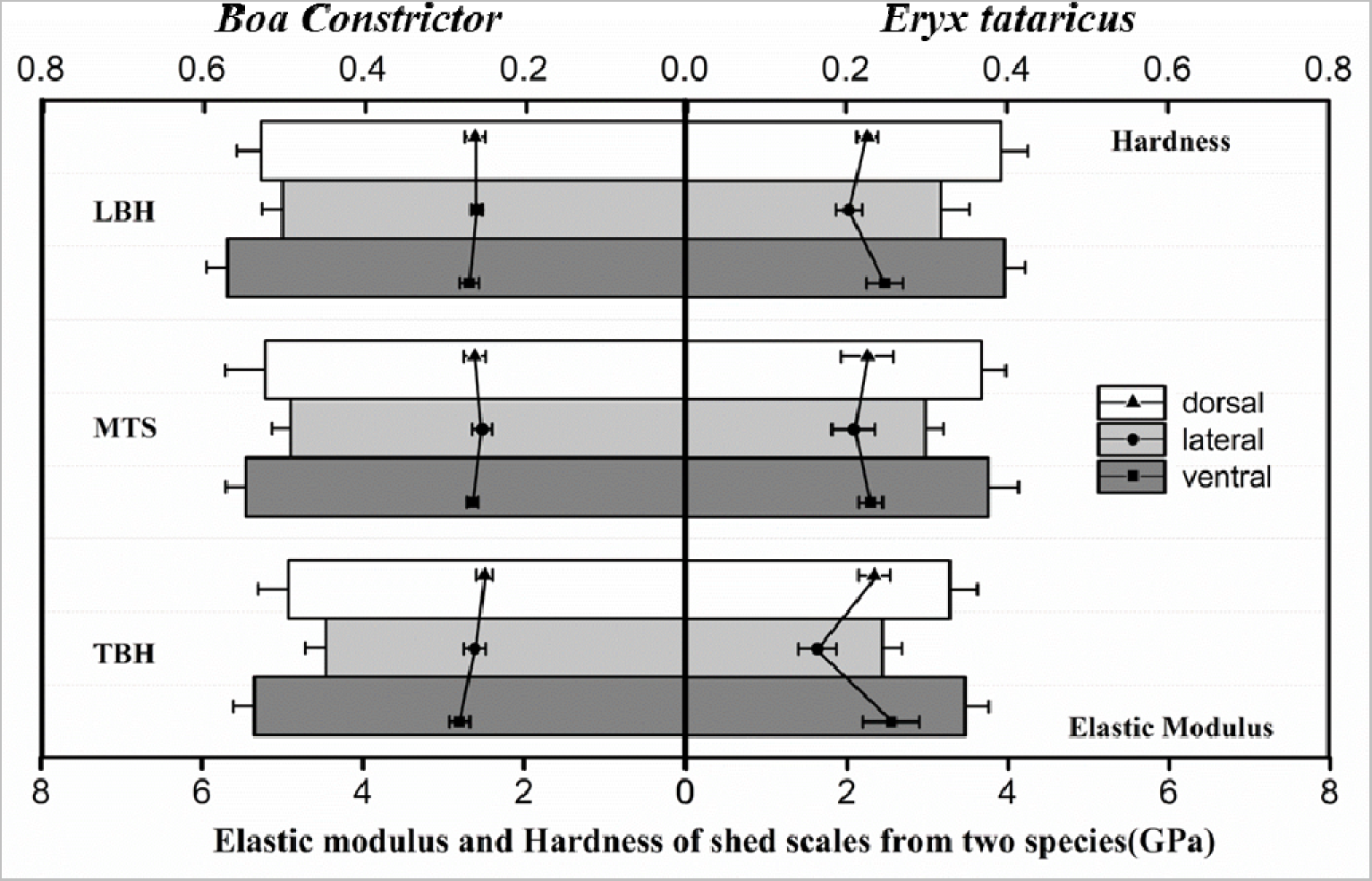
Elastic moduli (histograms) and hardnesses (lines) of scales at different sections of *Boa constrictor* and *Eryx tataricus*. The error bars denote standard deviations.

Pairwise statistical analyses (Students’ t test) have been performed on the elastic moduli and hardnesses of scales in different body positions (dorsal, lateral and ventral) and sections (LBH, MTS and TBH) for individual species (Fig. 8). The white boxes (diagonal from top left to bottom right in Fig. 8) show the significant differences among dorsal, lateral and ventral scales in the same body sections for *Boa constrictor* (Fig. 8A and 8B) and *Eryx tataricus* (Fig. 8C and 8D) in terms of elastic modulus (Fig. 8A and 8C) and hardness (Fig. 8B and 8D). It is found that both species exhibit a comparable pattern of elastic modulus differences with substantially lower elastic moduli for the lateral scales at distinct body sections (white boxes in Fig. 8A and 8C), whereas the differences for scale hardnesses from *Eryx tataricus* are more significant than those from *Boa constrictor*, especially at LBH (top left white boxes in Fig. 8B and 8D). The colored boxes (filled with orange, blue and green) display the significant differences among LBH, MTS and TBH at the same body positions for the two species. It is noted that for elastic modulus and hardness, body sections play a more important role for scales from *Eryx tataricus* than those from *Boa constrictor*, particularly for the comparisons of MTS vs. LBH and MTS vs. TBH (orange and blue boxes in Fig. 8). In summary, the mechanical properties of scales show dramatic variations for *Eryx tataricus* with regard to both body positions and sections, whereas body positions are more dominant in scale mechanical performance than body sections for *Boa constrictor*.

**Figure 8.**
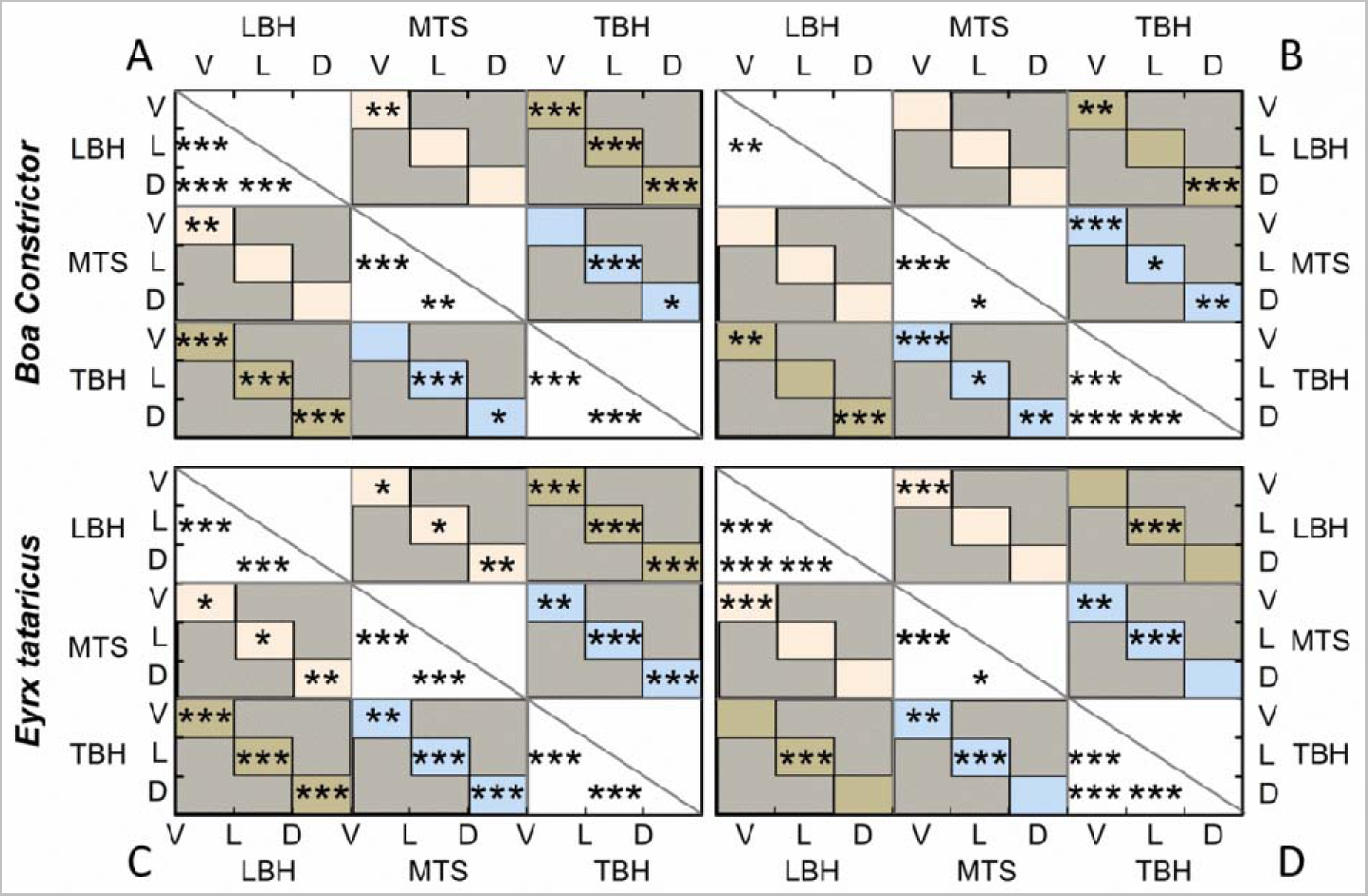
The significant differences of mechanical properties for scales from different body sections of *Boa constrictor* (A and B) and *Eryx tataricus* (C and D). (A) and (C) show the significant differences of elastic modulus for *Boa constrictor* and *Eryx tataricus*, respectively. (B) and (D) exhibit the significant differences of hardness for *Boa constrictor* and *Eryx tataricus*, respectively. Significance of Students’ t-test: * for p<0.05; ** for p< 0.01; *** for p<0.001). D: dorsal scales; L: lateral scales; V: ventral scales. Diagonal white boxes are the significant differences for scales at the same body section but different positions. Orange, blue and green boxes are the significant differences for scales at the same body position but different sections. The grey boxes are not applicable.

### 3.3 Tribological behaviors of scales

Anisotropic frictional properties have been demonstrated for scales of *Lampropeltis getula californiae* (Baum et al., 2014c). Herein, the friction coefficients of the scales from MTS of two species were measured in different rubbing directions by a nanotribometer to investigate the anisotropic frictional properties of scales in different body positions (Fig. 9). The friction coefficients were acquired in three directions: forward (F), backward (B) and transverse (T). Overall, the friction coefficients of scales from *Eryx tataricus* are larger than those of the counterparts from *Boa constrictor* (buildup boxes in Fig. 9A), particularly at dorsal and ventral positions. Moreover, the ventral scales show higher friction than other positions in general for each species. It is also apparent that for both species, the friction coefficients of the backward direction are higher than the other directions, indicating evident friction anisotropy.

**Figure 9.**
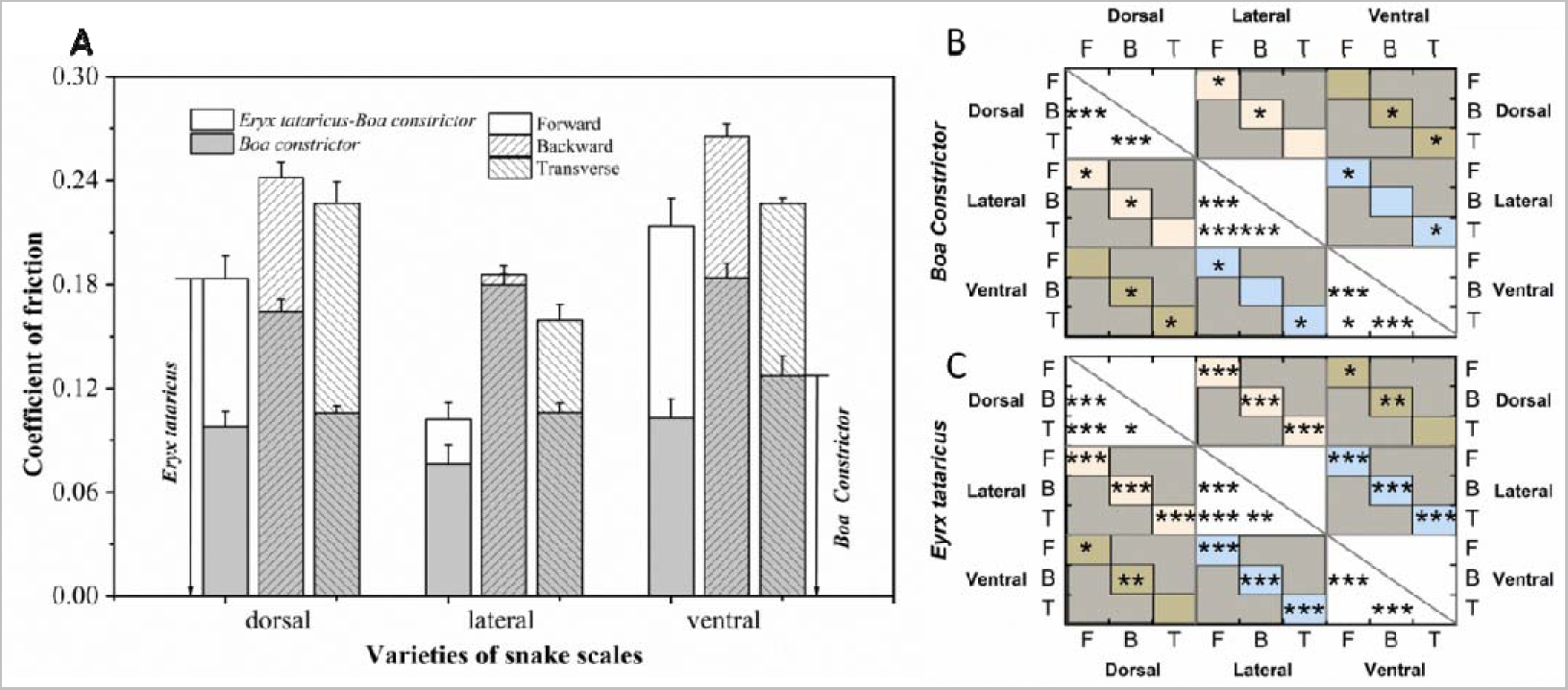
Anisotropic friction for scales in MTS of *Boa constrictor* and *Eryx tataricus*. (A) shows frictional coefficients of different rubbing directions for dorsal, lateral and ventral scales in MTS of *Boa constrictor* and *Eryx tataricus*. The error bars denote standard deviations. (B) and (C) are the significant differences of frictional coefficients in panel A. Significance of Students’ t-test: * for p<0.05; ** for p< 0.01; *** for p<0.001). F: forward; B: backward; T: Transverse. Diagonal white boxes are the significant differences for friction coefficients of the same body position but different rubbing directions. Orange, blue and green boxes are the significant differences for friction coefficients of the same rubbing direction but different body positions. The grey boxes are not applicable.

Pairwise statistical analyses (Students’ t test) have also been carried out on the friction coefficients in three directions for each species (Fig. 9B and 9C). The white boxes (diagonal from top left to bottom right in Fig. 9B and 9C) show the significant differences of friction coefficients among forward (F), backward (B) and transverse (T) directions at the same body positions for *Boa constrictor* (Fig. 9B) and *Eryx tataricus* (Fig. 9C). The colored boxes (filled with orange, blue and green) display the significant differences of friction coefficients with the same direction among dorsal, lateral and ventral scales for the two species. Interestingly, both species show a comparable pattern of friction coefficient differences at the same body positions (white boxes in Fig. 9B and 9C) indicating an evident anisotropic frictional performance, whereas the differences of the same rubbing direction at different body positions (dorsal, lateral and ventral) from *Eryx tataricus* are more significant than those from *Boa constrictor*, especially for dorsal vs. lateral and ventral vs. lateral (orange and blue boxes in Fig. 9B and 9C). In brief, the scales of the two species exhibit significant anisotropic frictional properties, while the differential friction of altered body positions is more prominent for *Eryx tataricus* than those for *Boa constrictor*.

## 4. Discussion

This work has investigated the morphology, mechanical properties and frictional behaviors of the scales from *Boa constrictor* and *Eryx tataricus*. Our morphological characterizations and nanoindentation measurements have corroboratively revealed substantial differences between the two species in terms of morphology, surface roughness and mechanical properties. Simultaneously, friction coefficients of scales in different body positions from the two snakes also exhibit significant anisotropy. Interestingly, the ventral scales show higher friction coefficients but lower surface roughness, together with relatively larger elastic modulus and hardness. Based on these results, we have proposed an assumption that the frictional performance of the scales, particularly some unexpected results, is owing to the coupling effect of the morphology and mechanical properties. This view is also supported by another recent work, which has hypothesized that the abrasive resistance of snake scales is not only due to a gradient in material properties of the scales, but also influenced by the morphology of scales (Klein and Gorb, 2014).

Both SEM and AFM images show that the morphologies on the scale surface are visibly different between the two species (Fig. 3 and Fig 5). The morphological divergence is likely a consequence of both respective inhabiting environments and different locomotion. *Boa constrictor* lives in a broad range of environments, which is less demanding in general compared to that of *Eryx tataricus* (Stidworthy, 1974). Thus, although the MTS of *Boa constrictor* takes more responsibility in its locomotion, the microstructure of ridges on the belly ventral scales is still evident. In contrast, the *Eryx tataricus* mainly resides in desserts and its ventral scales show an almost smooth surface of low roughness without clearly discernible micro-ornaments, probably caused by constant wearing on sand. Moreover, the whole body of *Eryx tataricus* is used in movement, especially in sand burrowing, which suggests that the dorsal and lateral scales also function in the propulsion. Hence, the morphology observed on the dorsal and lateral scales from *Eryx tataricus* is similar to those seen on the ventral scales from MTS of *Boa constrictor*, which can be rationalized by the reason to generate propulsion force.

More interestingly, the front edges of the scales of *Eryx tataricus* are mounted by tiny wrinkles resembling aeolian sand ripples. It is known that these sand ripples are resulted by repeated erosions of two-phase flow, or airflow, over a long time (Carter et al., 1980; Hutchings, 1977). Such a resemblance may not simply be a coincidence. Instead, it suggests that the morphology on the scales of *Eryx tataricus* could also be shaped by the factors that drive the formation of sand ripples. Furthermore, the patterns on the middle portions of dorsal and lateral scales from *Eryx tataricus* can be considered as the derivatives of the front-edge wrinkles, whereas the middle section of ventral scales only retain indiscernible microstructures due to more wearing. Together, the microstructures on the scale surface of *Eryx tataricus* may be an optimal formation adapting to the desert and sand-burrowing lifestyle. In contrast, there is no such a strong relationship between the morphology of the scales from *Boa constrictor*, assumingly due to its more complex surroundings and sophisticated lifestyle.

In terms of mechanical properties, both vertical and horizontal gradients are found on the scales of the two snakes (Fig. 6). The layout of repeating gradients on the whole snake body renders an integrated pattern of alternating hard-soft materials, which may afford an anti-wear function in their locomotion. The soft portions in the scales may not only extend and contract with muscle action during locomotion, but also dissipate the stress on the hard portions to reduce fatigue. Instead, the hard portions are equipped with higher elastic modulus to resist deformation as well as abrasive wear. Comparatively, the scales of *Boa constrictor* have larger elastic moduli and hardnesses than those of *Eryx tataricus*, which is likely to bear greater own body weight. Furthermore, the ventral scales in the same body sections of the two species are of the largest elastic moduli and hardnesses (Fig. 7), which may contribute to forward propulsion by increasing the friction during the undulation locomotion, as discussed below.

To our surprise, the investigation on the tribological behaviors of the scales from the two species has revealed higher friction coefficients for the ventral scales (Fig. 9A), which are of least surface roughness (Fig. 4) but relatively larger elastic modulus and hardness (Fig. 7). Intuitively, it is assumed that lower surface roughness with higher mechanical strength would result in less friction according to daily observations. However, this may not be applicable to the situation of microstructures. In our case of snake scales, the friction is influenced by both the roughness in morphology and the mechanical properties. In order to qualitatively illustrate the coupling effect of such two dimensions on friction, a 2x2 graph has been plotted assigning “+” and “-” signs to these factors based on the common beliefs of their impact on friction. For roughness, high roughness and low roughness are given “+” and “-” signs, respectively. For mechanical properties, high elastic modulus and hardness is considered as a benefit for low friction and thus given a “-” sign, whereas the opposite “+” sign represents low elastic modulus and hardness. In Fig. 10, double “+” or “-” would result in high friction due to a coupling effect of two factors. The “normal” coupling of high roughness (+) and low elastic modulus and hardness (+) is understandable, as seen with Velcro. On the contrary, the synergistic increase of friction due to the “double-crossed” coupling of low roughness (-) and high elastic modulus and hardness (-) is counterintuitive, with the mechanism being elusive. One possible explanation is that the undeformable surface can retain more stress on the friction interface, which affords more driving force for molecular interaction. Meanwhile, the smooth morphology is able to maximize the real contact area of the friction to engage more molecules for interaction. Hence, in comparison with the lateral scales, the ventral scales of the two species, although with lower surface roughness and larger elastic modulus and hardness, have demonstrated higher friction coefficients. Such a coupling effect on ventral scales may also be an optimized adaptation to sliding locomotion, which not only smoothens the ventral surface, but also demands more load-bearing capability at the belly.

**Figure 10.**
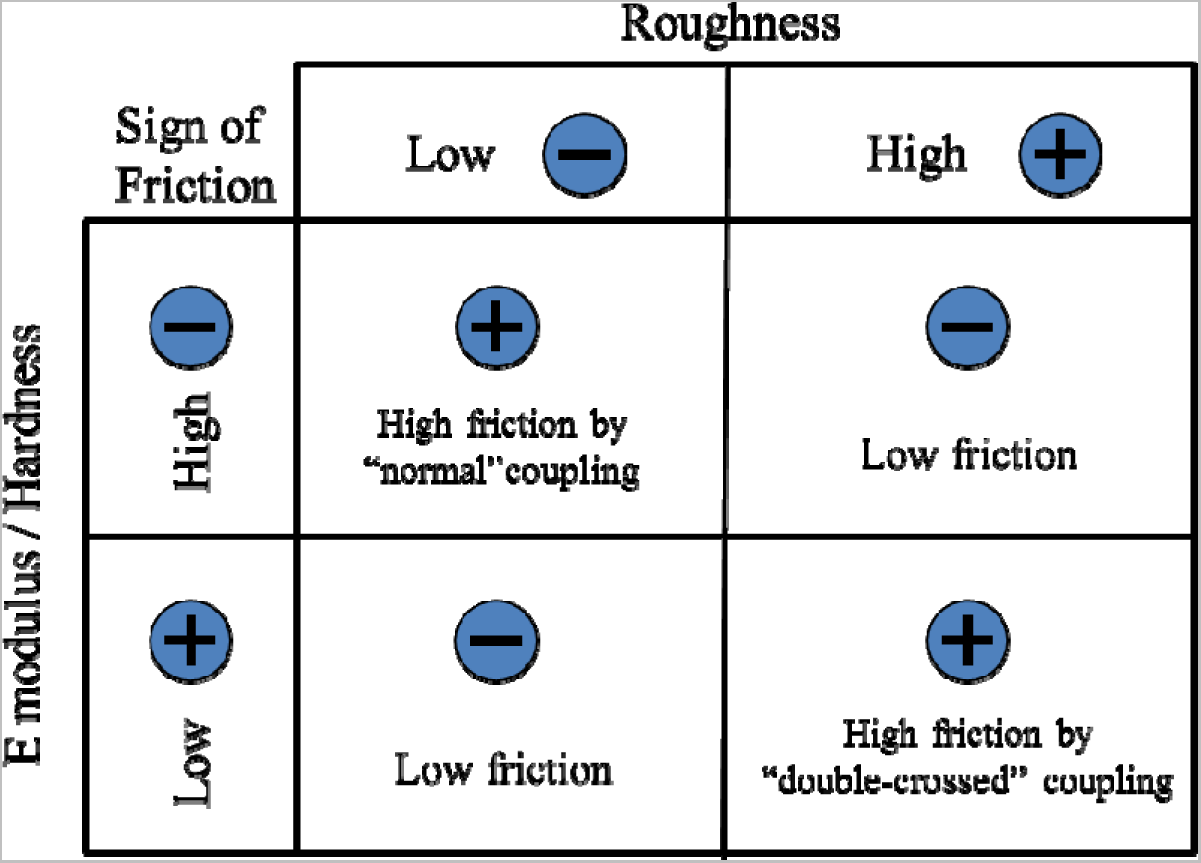
The illustration for the “normal” and “double-crossed” coupling mechanism of morphology and mechanical properties on frictional behaviors.

In addition, our tribological characterization has also revealed obvious anisotropic friction for the two snakes (Fig. 9A), in line with previous studies (Hu et al., 2009). Based on previous work, it is assumed that the ridged microstructures provide propulsion force for snakes, almost perpendicular to the direction of locomotion. Moreover, the patterns on the scale surfaces incur an interlocking effect with the irregularities and asperities of the ground, which leads to high friction to prevent slithering in backward direction (Alexander, 2003). Thus, the anisotropic frictional behavior of snake scales is a consequence of evolution adapting to locomotion and lifestyles.

## 5. Conclusion

In this work, the morphology and mechanical properties of the scales from different body sections and positions of *Boa constrictor* and *Eryx tataricus* have been characterized and compared to investigate the corresponding effects on the tribological behaviors and to probe the possible coupling mechanism. The morphological characterizations of SEM and AFM have revealed significant differences between the two species with the roughness of scales from *Boa constrictor* being larger in general. The mechanical properties measured by nanoindentation have corroboratively demonstrated substantial differences in terms of elastic modulus and hardness. Meanwhile, tribological characterizations of scales in different body positions from the two species also exhibit evident anisotropy. Interestingly, the ventral scales manifest higher friction coefficients but lower surface roughness, together with relatively larger elastic modulus and hardness. A “double-crossed” hypothesis has been proposed to explain the observed coupling effect of the morphology and mechanical properties on friction, which may afford valuable insights for the design of materials with desirable tribological performance.

## Acknowledgements

This work was supported by National Natural Science Foundation of China (51375204 and U1601203) and Jilin Provincial Science & Technology Department (20140101056JC). The authors thank Dr. Zhonghao Jiang from College of Materials Science and Engineering, Jilin University, for the help on nanoindentation tests and Dr. Zhanjiang Yu from Changchun University of Science and Technology for the help on tribological measurements.

